# A multimodal dataset of EEG, eye-tracking, and physiological signals during naturalistic smartphone interactions

**DOI:** 10.64898/2026.04.21.719334

**Authors:** Prakash Mishra, Tapan K. Gandhi, Saurabh R. Gandhi

## Abstract

Smartphones have become pervasive tools for communication, information consumption, and digital interaction, yet the neurophysiological dynamics associated with naturalistic smartphone use remain insufficiently characterized. Here, we present a multimodal physiological dataset collected during ecologically valid smartphone interaction and a subsequent relaxation condition. Twenty-three participants engaged with their most frequently used smartphone application (primarily gaming or short form video) for ten minutes, followed by a five-minute passive viewing of a standardized nature video. Simultaneous recordings were obtained from electroencephalography (EEG; 64 channels), wearable eye-tracking, photoplethysmography (PPG), and galvanic skin response (GSR) sensors. Questionnaire-based assessments, including the smartphone addiction scale (SAS) and the mobile phone problematic use scale (MPPUS), are also collected to characterize individual differences in smartphone-related behavioral traits. All data streams are synchronized using transistor-transistor logic (TTL) trigger signals to ensure precise temporal alignment across modalities. The dataset is organized according to the Brain Imaging Data Structure (BIDS) specification and is publicly available on OpenNeuro.org (Accession Number: ds007537). This dataset enables the investigation of neural, ocular, and autonomic responses during smartphone interaction and supports multimodal analysis of diverse smartphone behaviors while preserving ecological validity.

## Background and summary

Being a part of day-to-day life, smartphones have become ubiquitous cognitive extensions for billions of individuals worldwide^1^. Smartphone usage is driven by the type of application used by individuals, supporting a wide range of activities beyond traditional communication, including social media interaction, short-form video consumption, gaming, and information browsing^2–4^. Behavioral research has extensively examined the cognitive and psychological effects of smartphone use, demonstrating associations with attentional modulation, altered task-switching behavior, and changes in emotional regulation^5–7^. Studies have shown that frequent smartphone interaction may influence sustained attention, increase susceptibility to distraction, and modify habitual information-seeking behavior^8,9^. In addition, behavioral assessments have linked patterns of smartphone use with sleep disruption, reduced cognitive performance in attention-demanding tasks, and altered patterns of engagement during academic and occupational activities^2,10–12^.

Different categories of smartphone activities impose distinct perceptual and cognitive demands, resulting in activity-specific physiological and behavioral responses^4^. Short-form video applications, for example, present rapidly changing visual stimuli that impair sustained attention and promote frequent attentional switching, ultimately contributing to heightened distractibility and reduced concentration.^13,14^. Gaming applications often require sustained visuomotor coordination, rapid decision-making, and continuous performance monitoring, which can increase cognitive workload and attentional demand^15,16^. Healthy, moderate gaming acts as beneficial engagement that trains sensorimotor responsiveness and visual attention^15^. Conversely, pathological gaming (e.g., internet gaming disorder) transitions this workload into chronic overload, demanding excessive, compensatory cognitive resources from executive networks to maintain basic inhibitory control^16^.These differences suggest that smartphone interaction represents a heterogeneous behavioral context, in which the type and structure of digital content influence neural and physiological responses.

In addition to behavioral assessments, physiological measures have been increasingly used to investigate the neural and autonomic correlations of smartphone use. EEG has been employed to characterize neural activity associated with attentional engagement, cognitive workload, and functional connectivity during digital interaction^17,18^. Prior EEG studies have documented oscillatory modulations under well-defined digital task conditions. Alpha-band (8-13 Hz) desynchronization has been observed during periods of sustained visual attention and active information processing on screen-based devices, reflecting engagement of top-down attentional networks^19,20^. Frontal theta-band (4–8 Hz) power increases have been reported during conditions of elevated cognitive load, including working memory-intensive tasks such as active text composition and multitasking on mobile devices^21,22^, and beta-band (13-30 Hz) suppression has been linked to motor preparation and execution during touchscreen input^23^. Eye-tracking studies have further demonstrated that smartphone interaction such as gaming on phone, influences gaze dynamics, fixation patterns, and visual attention allocation, reflecting underlying perceptual and attentional mechanisms^24–26^. Pupillary dilation, a well-established proxy for cognitive load mediated by the locus coeruleus-noradrenergic system, has been shown to scale with the cognitive demands of smartphone tasks^27^. Peripheral physiological measures, such as cardiac and electrodermal activity, have also been used to assess autonomic nervous system responses during technology use, providing indicators of physiological arousal and regulatory processes^28,29^. Smartphone use under different contexts has thus been shown to reflect in physiological changes in EEG, pupillometry, gaze, cardiac and electrodermal activity.

These efforts have established a strong foundation for understanding specific aspects of smartphone-related neurophysiological responses. Crucially though, cross-modal relationships, such as the coupling between frontal theta power and pupillary dilation under working memory load^27,30^, or the co-modulation of alpha desynchronization and fixation stability during attentional engagement^19,20^, represent emergent psychophysiological phenomena that can only be observed when modalities are acquired concurrently and analyzed jointly. Independent unimodal recordings preclude this investigation due to unavoidable inter-session and inter-individual variability that would confound any post-hoc attempt at integration. The simultaneous acquisition of these complementary modalities enables an integrated characterization of psychophysiological responses during technology interaction and builds upon recent multimodal approaches for assessing user engagement and cognitive processing in ecological digital interaction settings^31,32^.

The majority of prior EEG and physiological studies of smartphone behavior have employed experimenter-assigned tasks such as reading standardized passages, viewing preselected videos, or following prescribed typing prompts to ensure stimulus control and interparticipant comparability^21,33^. While such designs afford experimental control, they may not capture the neural and physiological dynamics characteristic of spontaneous, self-directed smartphone use, in which content selection, interaction pacing, and motivational engagement differ markedly from imposed laboratory tasks^34^. Building upon this foundation, the development of open-access datasets integrating multiple physiological modalities, incorporating participants’ naturally preferred smartphone applications, would further advance the field.

The growing interest in the neurophysiology of smartphone behavior has underscored the need for comprehensive datasets that integrate self-reported questionnaire-based addiction profiling with simultaneous multimodal physiological recordings during ecologically valid smartphone use^35–37^. To support research in this area, we developed a multimodal physiological dataset acquired during ecologically valid smartphone interaction and a subsequent standardized relaxation condition. The experimental paradigm consisted of two sequential phases. In the first phase, participants interacted freely with their self-selected preferred smartphone application for a duration of 10 minutes, enabling the capture of physiological responses during naturalistic and self-directed smartphone use, primarily gaming or short form video. In the second phase, participants viewed a standardized, relaxing 5-minute nature video, providing a controlled condition for assessing physiological activity during reduced cognitive engagement. Physiological signals were recorded simultaneously across modalities, ensuring precise temporal synchronization.

The dataset includes recordings from 30 participants, each consisting of synchronized multimodal 64 - channel EEG, wearable eye-tracking, PPG, and GSR signals along with the smartphone usages based psychometric questionnaire responses. In addition, the experimental design includes both smartphone interaction and a standardized passive-viewing comparison condition, enabling within-subject comparisons. The combination of multiple physiological modalities and synchronized recordings provides a rich dataset for multimodal analysis. The structured and open-access organization of the dataset enables reproducible analysis and supports a wide range of applications, including physiological signal analysis, multimodal integration, cognitive state modeling, and machine learning-based classification

## Methods

### Ethics statement

The experimental protocol and data sharing is approved by the “Indian Institute of Technology, Delhi (IIT-Delhi) Institutional Human Ethics Committee” approval number VS-1225-REA101. Participants provided explicit consent to share anonymized data on open access platforms. Participants were briefed about the experimental procedure and the equipment used before the start of the recording session.

### Participants

Thirty participants (10 females) voluntarily participated in the experiment. Participants are recruited through an online Google Form shared with students by word of mouth and are between 20 and 33 years of age (mean age = 25.1years). All the participants have normal or corrected to normal vision with no history of neurological disorders.

### Experiment Protocol

Participants are seated in a comfortable chair and asked to complete the informed consent form. Following the consent procedure, the participant undergoes the experimental setup for physiological sensing devices (64-channel EEG, wearable eye-tracking glasses and PPG and GSR sensors), lasting around 45 minutes. Participants are allowed to use their smartphones during this preparation phase, which allows them to acclimatize to the laboratory environment before the actual experimental session begins. The experiment is conducted in a small, quiet and non-acoustically treated room with only the participant and experimenter present with minimal or no surrounding noise.

The experimental paradigm consists of two sequential conditions. In the first one (naturalistic smartphone use, lasting 10 mins), participants use their personal smartphones and are instructed to open and freely interact with one of their most frequently used applications. Participants remain seated comfortably and hold the smartphone in a natural viewing position during this condition. Participants are free to interact with the smartphone using their left hand, right hand, or both hands according to their usual interaction style. The selected hand-use condition remains consistent throughout the smartphone-use session and is recorded by the experimenter. The physiological sensor placement on the left hand leaves the thumb and index finger available for touchscreen interaction and therefore does not preclude two-handed smartphone interaction. No constraints are imposed on the interaction style, allowing participants to engage freely with the application in their typical manner and device setting (e.g. screen size, resolution, brightness) except for minimizing body movements other than finger movements required for smartphone interaction (Figure 1). Participants use audio through the smartphone’s built-in speaker, with the volume controlled according to their usual preference. No headphones or earpieces are used. The experimenter records whether audio is used by each participant during this task. The application is selected by the participant based on their usual smartphone-use behaviour and is not assigned or restricted by the experimenter, so as to preserve ecological validity and capture naturalistic smartphone interaction behavior. The type of application used is recorded by the experimenter. Only the broad activity category is retained in the released dataset to protect participant privacy. The selected applications include short-form video platforms (n = 15; e.g., Instagram Reels, YouTube Shorts), gaming applications (n = 13; e.g., Subway Surfers, Battlefield), and reading applications (n = 2; e.g., article reading platforms).

**Figure 1.**
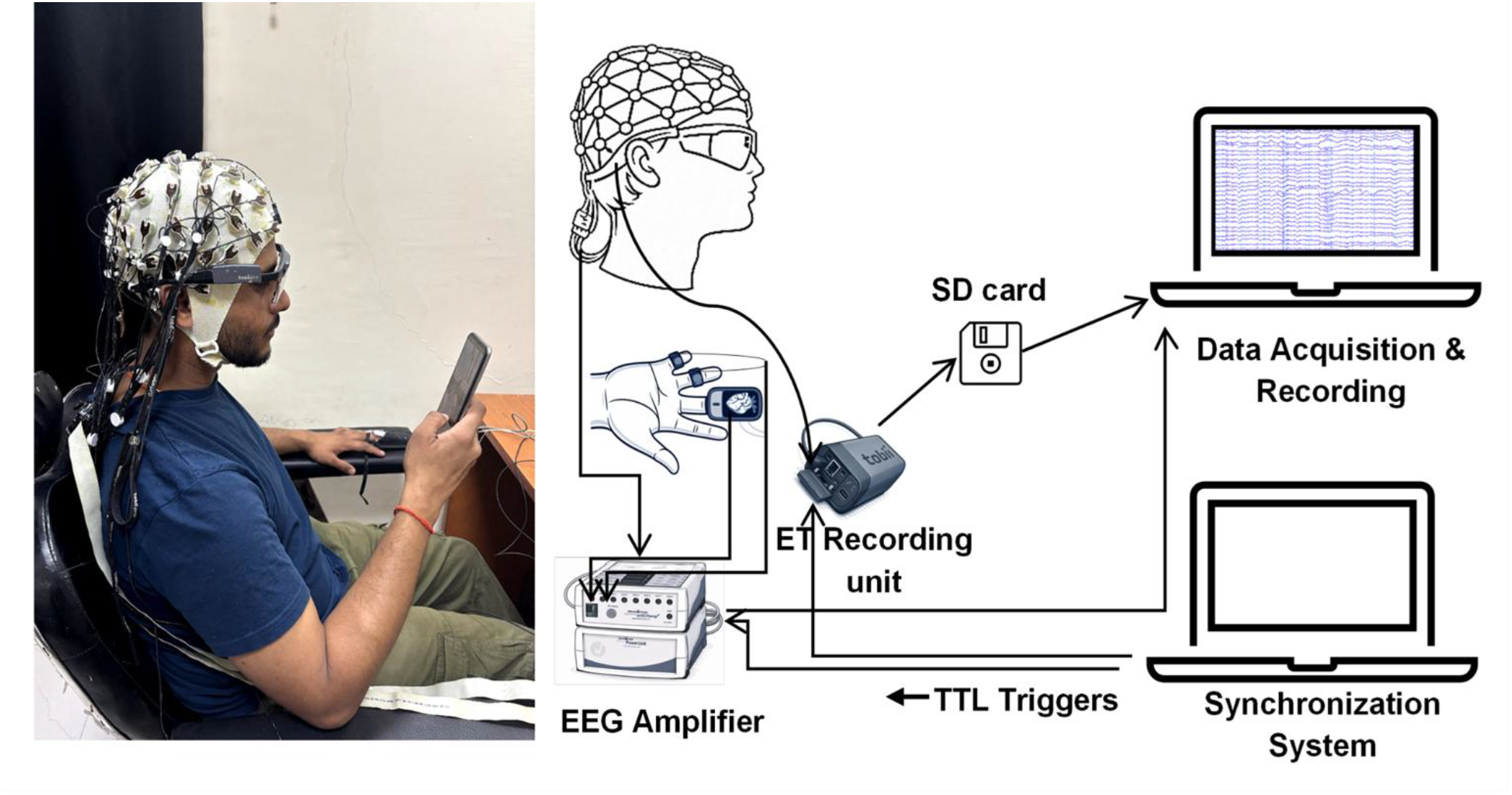
Experimental setup and system architecture for synchronized EEG, PPG, EDA and eye-tracking data acquisition. Participant wearing EEG cap and eye-tracking device during the experiment (Left) (illustrative image of one of the authors with permission). System diagram showing EEG amplifier, eye-tracking recording unit, data acquisition computer, and synchronization system connected via TTL triggers (Right).

This naturalistic smartphone use condition is followed by the second condition (relaxing nature video, lasting 5 mins). In this phase, participants place their smartphones face-down and view a standardized high-definition nature video (E*ARTH 4K-Relaxation Film-Peaceful Relaxing Music-Nature 4K Video UltraHD-OUR PLANET from youtube*.*com);* segment from 08:00 to 13:02 minutes) presented on a computer monitor. The monitor is positioned at a viewing distance of approximately 85 cm from the participant. Participants are instructed to watch the video while remaining seated. Participants listen to the audio through stereo desktop speakers for this task, with the volume maintained at a comfortable and consistent level across participants.

After completion of both experimental sessions, participants complete the original 33-item Smartphone Addiction Scale (SAS)^36^ and the original 27-item Mobile Phone Problematic Use Scale (MPPUS)^35^ using a Google Form. The SAS uses a six-point Likert response scale, whereas the MPPUS uses a ten-point Likert response scale. Total scores are computed by summing the individual item responses, with higher scores indicating greater levels of smartphone addiction and problematic mobile phone use, respectively. Participants complete the original English versions of both questionnaires.

### Recording Equipment and Signal Acquisition

EEG signals are recorded using the Brain Products ActiCHamp 64-channel system (Brain Products GmbH) equipped with active Ag/AgCl electrodes positioned according to the international 10-20 system. The EEG data are acquired at a sampling rate of 1000 Hz with Cz used as the reference electrode during recording. Peripheral physiological signals are recorded using auxiliary analog inputs (AUX1 and AUX2) of the ActiCHamp amplifier. The PPG signal is recorded using the Brain Products Blood Pulse Sensor connected to AUX1 and attached to the middle finger of the participant’s left hand. The GSR signal is recorded using the Brain Products GSR Sensor connected to AUX2 and attached to the ring and little fingers of the participant’s left hand. Both signals are sampled synchronously with the EEG at 1000 Hz. The GSR channel is configured in the BrainVision Recorder workspace with a gradient of 25 and an offset of 0. The Blood Pulse Sensor outputs a relative photoplethysmographic waveform and therefore does not provide an absolute physical calibration. Eye-tracking data are collected using the wearable Tobii Pro Glasses 2 eye tracker. The system is calibrated individually for each participant prior to the experiment and records gaze position, pupil diameter, gyroscope, and accelerometer data during the experimental sessions.

Synchronization between the ActiCHamp EEG system and the Tobii Pro Glasses 2 eye tracker is achieved using TTL trigger signals. TTL pulses are transmitted simultaneously to the Tobii Pro Glasses 2 via its external synchronization interface and to the ActiCHamp system through its standard digital trigger input port. At the beginning and end of each experimental session, a sequence of five TTL trigger pulses is delivered at 1-second intervals to mark the session boundaries. During the experimental session, additional periodic TTL trigger pulses are generated at fixed intervals of 20 seconds to maintain synchronization and provide continuous temporal reference points throughout the recording (Figure 2). These trigger events are recorded concurrently in both the eye-tracking data stream and the EEG acquisition system, providing precise temporal markers for the alignment of multimodal recordings.

**Figure 2.**
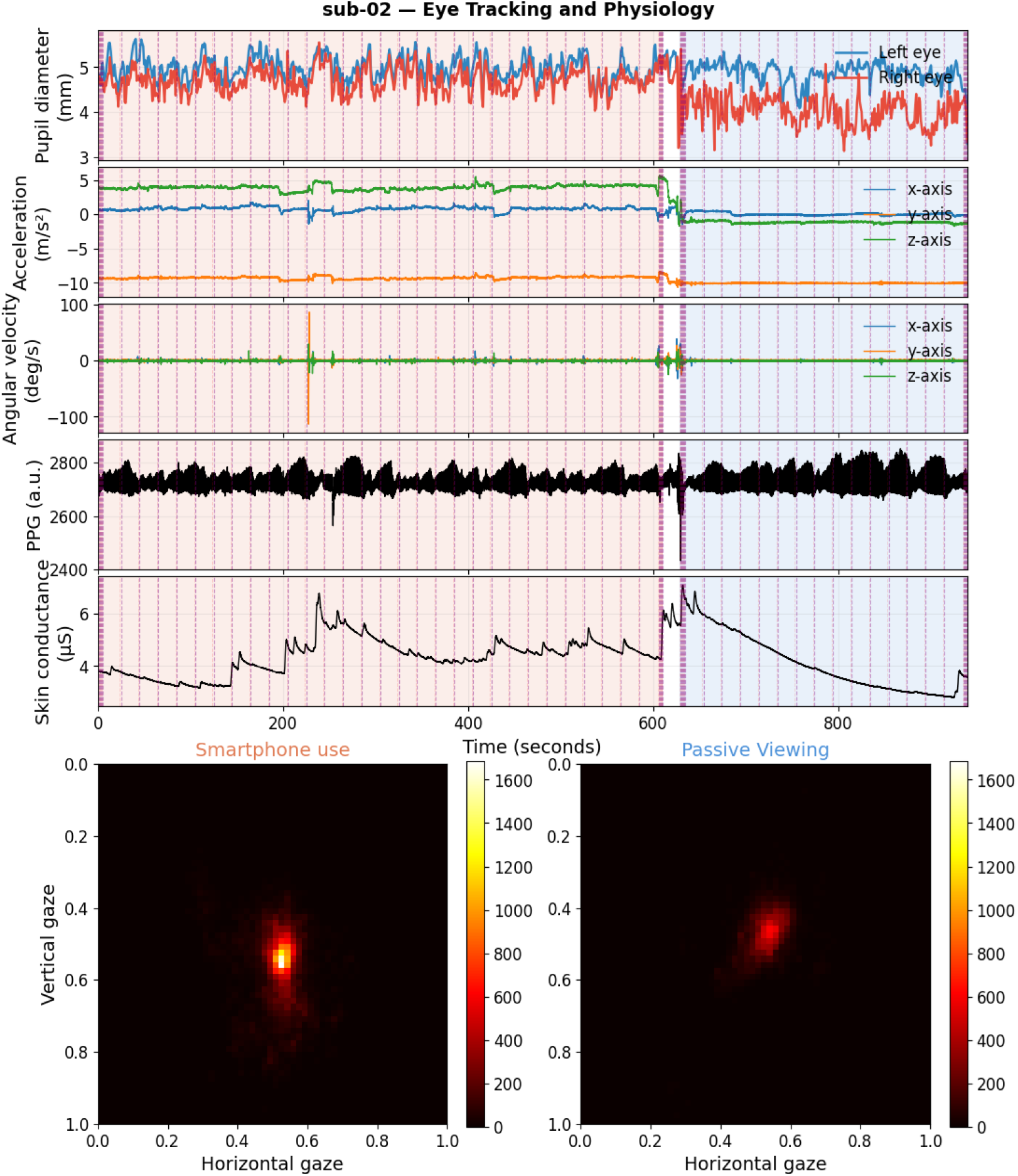
Multimodal physiological and eye-tracking responses during Smartphone Use (SM) and Passive-viewing (PV) sessions. Time-series plots show pupil diameter (left and right eyes), head acceleration, angular velocity, photoplethysmography (PPG), and skin conductance (GSR), synchronized on a common time axis. Shaded regions indicate task periods identified using TTL pulse clusters: Smartphone use (orange) and Passive-viewing (blue). Vertical dashed lines represent TTL trigger pulses, and purple lines indicate experimental event markers. The lower panels show two-dimensional gaze density heatmaps for each session, illustrating spatial gaze distribution normalized to the display coordinate system. Warmer colors indicate higher gaze density. These results demonstrate distinct physiological and oculomotor patterns across task conditions.

### Data Records

The dataset is available at OpenNeuro (reference number ds007537.v2.0.2 ^38^) and is organized according to the Brain Imaging Data Structure (BIDS) specification (Figure 3). The raw dataset adheres to the standard BIDS (validator v 3.0.1) directory hierarchy, ensuring compatibility with commonly used neuroimaging and physiological data analysis tools.

**Figure 3:**
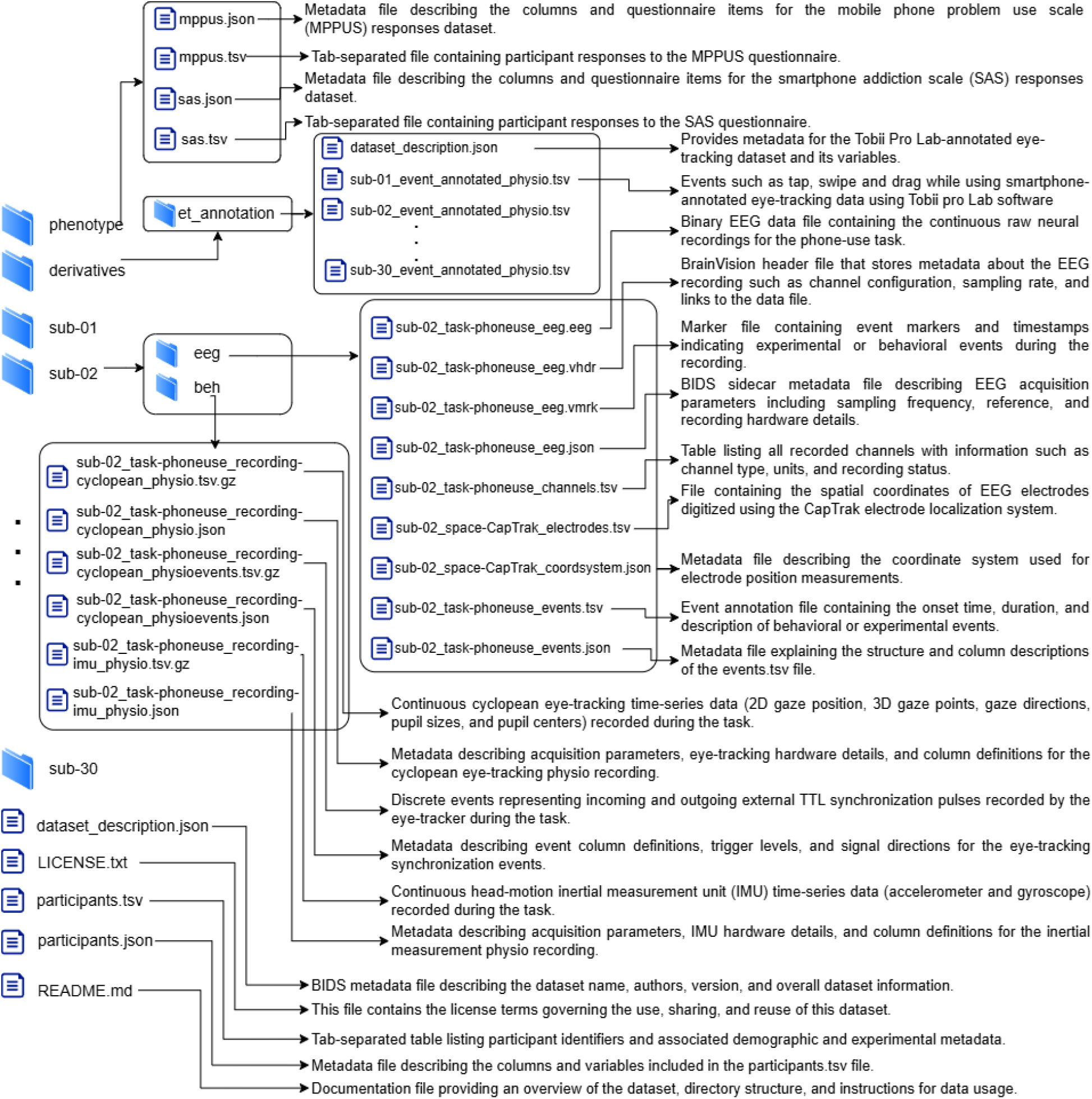
Directory structure of the multimodal smartphone interaction dataset organized according to the BIDS specification. The dataset includes questionnaire responses (MPPUS and SAS), eye-tracking recordings, and EEG data collected during the smartphone-use task. EEG recordings are stored in BrainVision format (.eeg, .vhdr, .vmrk) with accompanying metadata (.json), channel information (channels.tsv), electrode locations digitized with CapTrak (electrodes.tsv, coordsystem.json), and event annotations (events.tsv, events.json). Eye-tracking data are provided in tabular format with corresponding metadata files, while questionnaire responses are stored in TSV files with accompanying JSON descriptions.

At the top level, the dataset includes the files *dataset_description*.*json, README*.*md, participants*.*tsv*, and *participants*.*json. The dataset_description*.*json* file provides general information about the dataset, including the dataset name, authors, and version. The README.md file contains documentation describing the dataset organization and usage instructions. Participant metadata are stored in *participants*.*tsv*, and the corresponding metadata descriptions are provided in *participants*.*json*. Participant metadata include participant identifier (*participant_id*), age, biological sex, smartphone application type (*app_type*), hand used during smartphone interaction (*hand*), audio usage (*audio*), and smartphone screen orientation (*orientation*) during the smartphone session.

In addition to physiological recordings, the dataset includes questionnaire responses assessing smartphone usage behaviors. The *phenotype* directory contains behavioral questionnaire data assessing smartphone use-related behavioral characteristics. These data are provided in tab-separated value (.tsv) format following BIDS conventions. The files *sas*.*tsv* and *mppus*.*tsv* contain participant responses to the SAS and the MPPUS respectively. Metadata describing the questionnaire items and column definitions are provided in the corresponding .*json* files.

Each participant is assigned a unique identifier and stored in a dedicated directory labeled sub-XX, where XX denotes the participant ID. Within each participant directory, modality-specific subdirectories contain the corresponding data files. The EEG recordings are stored in the *eeg* directory and follow the BrainVision file format. Each recording includes the raw EEG signal file (*\*_eeg*.*eeg*), the corresponding header file (*\*_eeg*.*vhdr*), and the marker file (*\*_eeg*.*vmrk*) containing event markers recorded during the experiment. Additional metadata describing the acquisition parameters are provided in the sidecar file (*\*_eeg*.*json*). Channel information is stored in \**_channels*.*tsv*, which lists channel names, types, units, and recording status. Electrode positions digitized using the CapTrak system are provided in \**_space-CapTrak_electrodes*.*tsv*, while the coordinate system definition is stored in \**_space-CapTrak_coordsystem*.*json*. These files allow for reconstruction of the spatial configuration of EEG electrodes. Behavioral event annotations are stored in *\*_events*.*tsv*, which includes the onset time, duration, and description of events occurring during the smartphone-use task (recording onset and offset). The corresponding metadata describing the event columns are provided in *\*_events*.*json*. The PPG and GSR signals are stored together with the EEG data. Within the BIDS dataset, these auxiliary physiological channels are labelled PPG and GSR and documented in the accompanying \**_channels*.*tsv* metadata. The PPG signal is provided in arbitrary units (a.u.), whereas the GSR signal is reported in microsiemens (µS). The dataset was converted to the BIDS format using the MNE-BIDS library (version 0.16.0).

Eye-tracking data are acquired using the Tobii Pro Glasses 2 system, which stores raw eye-tracking data on the device’s secure digital (SD) card in *livedata*.*json*.*gz* format. A custom Python-based parser is used to convert the raw data into a BIDS-specific format. For each participant, eye-tracking recordings are stored within the subject-specific directory (sub-XX) within the *beh/* directory using BIDS-compliant naming conventions, where the *task-phoneuse* entity denotes the experimental condition involving naturalistic smartphone use. Continuous eye-tracking data (including cyclopean 2D gaze coordinates, 3D gaze points, gaze direction unit vectors, pupil sizes, and pupil center positions) are provided in compressed tab-separated value format (*\*_recording-cyclopean_physio*.*tsv*.*gz*) accompanied by a corresponding JSON sidecar file (*\*_recording-cyclopean_physio*.*json*), with *PhysioType* set to eyetrack. The first three columns of the gaze data file correspond to the required BIDS fields: *timestamp, x_coordinate, and y_coordinate*, representing the temporal index and gaze position of the recorded eye. Inertial measurement unit (IMU) data (containing 3-axis accelerometer and gyroscope recordings) are stored separately in *\*_recording-imu_physio*.*tsv*.*gz* accompanied by *\*_recording-imu_physio*.*json*, with PhysioType set to generic. External TTL synchronization signals used to align the eye-tracking and EEG timelines are recorded in a separate events file *(*_recording-cyclopean_physioevents*.*tsv*.*gz)* and JSON sidecar, containing columns for *onset, duration, trial_type, direction*, and *value*.

The JSON sidecar files provide essential metadata, including sampling frequency (*SamplingFrequency*), recording start time (*StartTime*), recorded eye (*RecordedEye*), and an ordered list of columns (*Columns*), along with detailed descriptions and units for each variable. TTL synchronization signals are included as additional columns in the eye tracking data file, enabling precise temporal alignment across modalities.

In addition to the raw recordings, the dataset includes annotated and exported eye-tracking data generated using Tobii Pro Lab (Version 26.10), located in the *derivatives/et_annotation/* directory. This directory contains a *dataset_description*.*json* file and a subject-specific *sub-xx_event_annotated_physio*.*tsv* file for each participant. The *dataset_description*.*json* file provides general information about the annotated eye-tracking dataset and includes descriptions of the variables contained in the *sub-xx_event_annotated_physio*.*tsv* files.

### Technical validation

#### EEG Signal Quality and Spectral Feature

EEG signal quality is assessed using an automated preprocessing and quality-control pipeline implemented in MNE (version 1.7.0)-Python (version 3.12.3) and applied to the BIDS-formatted dataset. The validation procedure aims to reduce artifacts, identify unreliable sensors, and quantify signal quality using spectral metrics. To reduce electrical interference, a notch filter is applied at the local power-line frequency (50 Hz). The signals are then band-pass filtered between 0.5 and 95 Hz using a second-order Butterworth filter with zero-phase (forward-backward) filtering. It preserve physiologically relevant EEG frequencies while retaining high-frequency components required for noise-floor estimation. Ocular artifacts are attenuated using Independent Component Analysis (ICA). Independent components are estimated from the filtered EEG signals, and components associated with eye movements are automatically identified using correlation-based detection (MNE-Python’s find_bads_eog method). In this method, each of six frontal electrodes (Fp1, Fp2, AF7, AF8, F7, F8) serves as an electrooculographic proxy, band-pass filtered at 1-10 Hz, and the Pearson correlation between every independent component and each proxy channel is computed separately. Correlation scores are standardised across components, and a component is flagged when its absolute z-scored correlation with any proxy channel exceeds 3.0 for any one of the six proxy channels. Up to five flagged components per participant (mean 4.57, range 2-5) are excluded before reconstruction. Component identification is fully automated to ensure reproducibility. To detect unreliable sensors, an automated bad channel detection procedure is applied using two criteria. First, channels exhibiting unusually high variance relative to other electrodes are flagged using the conventional ±3 SD outlier rule (variance z-score > 3.0^39^). Second, spatial consistency is evaluated by computing the Pearson correlation between each channel and its six nearest neighboring electrodes; channels with a mean correlation below 0.35 are flagged as potentially noisy. This threshold, comparable to the 0.4 criterion used in the PREP^40^ pipeline, was selected empirically from the distribution of neighbour correlations to flag clearly deviant channels. On average, 9.60 channels are flagged per participant (SD = 4.98; range 2-20). Identified bad channels are interpolated using spherical-spline interpolation, and the signals are subsequently re-referenced to the common average. For signal quality assessment, continuous EEG recordings are segmented into 20-s intervals aligned with experimental triggers corresponding to the smartphone-use and passive-viewing conditions. Segments containing large amplitude artifacts exceeding ±300 μV in any channel are automatically rejected. On average, 2.8 segments (4.7%) were rejected out of 60 in the SM session, and 1.9 segments (6.4%) were rejected out of 30 in the PV session per participant. The remaining artifact-reduced segments are concatenated separately for each condition.

Signal quality is quantified using a spectral signal-to-noise ratio (SNR) derived from the power spectral density estimated using Welch’s method with 4-s windows. Frequency-band SNR is computed for canonical EEG frequency bands (delta: 0.5-4 Hz, theta: 4-8 Hz, alpha: 8-13 Hz, beta: 13-30 Hz, and gamma: 30-45 Hz) as the ratio between the mean band power and a noise floor estimated from the 45-95 Hz frequency range, excluding frequencies within ±2 Hz of the power-line frequency. This high-frequency range (45-95 Hz) is dominated by non-neural contributions such as muscle activity, and environmental interference, and therefore provides a suitable proxy for broadband noise. By expressing SNR in decibels, this metric enables robust comparison across subjects and conditions. Higher SNR values indicate cleaner recordings with stronger physiological signals relative to noise, whereas lower values reflect increased contamination or reduced signal quality. Across the cohort, the SNR followed the expected spectral hierarchy, with delta exhibiting the highest SNR values for the large majority of participants (29 of 30; except sub-04), followed by theta and alpha, while beta and gamma showed lower SNR values (Figure 4A). This pattern is consistent with the characteristic 1/f spectral profile of scalp EEG, where lower-frequency oscillations have higher power relative to the noise floor. The group-level SNR hierarchy during smartphone usage is delta (14.31 dB) > theta (9.58 dB) > alpha (8.44 dB) > beta (5.22 dB) > gamma (3.18 dB), with positive mean SNR confirming that oscillatory power above the high-frequency noise floor is detected in every canonical band (Table 1). Whereas the hierarchy shifts to delta (15.29 dB) > alpha (10.91 dB) > theta (10.29 dB) > beta (6.08 dB) > gamma (3.07 dB) during the passive-viewing session. This condition-dependent shift in the alpha-theta ordering is a positive internal validity reflecting a known neurophysiological effect rather than a noise-driven artefact ^41,42^.

**Table 1.** Group-level spectral signal-to-noise ratio (SNR, dB) across canonical EEG frequency bands for the Smartphone and Video Viewing conditions (N = 30). Values represent mean SNR across subjects.

| **Band** | **Condition** | **Mean (dB)** | **SD (dB)** | **Median (dB)** | **Range (dB)** |
|---|---|---|---|---|---|
| **Delta**<br>**(0.5 - 4 Hz)** | Smartphone | 14.30 | 3.6 | 14.28 | 7.27 - 25.90 |
|  | Passive Viewing | 15.28 | 3.71 | 15.29 | 7.27 - 25.15 |
| **Theta**<br>**(4 - 8 Hz)** | Smartphone | 9.57 | 3.30 | 9.67 | 2.40 - 15.93 |
|  | Passive Viewing | 10.28 | 2.87 | 10.14. | 2.30 - 15.64 |
| **Alpha**<br>**(8 - 13 Hz)** | Smartphone | 8.44 | 3.56 | 8.13 | 0.26 - 16.14 |
|  | Passive Viewing | 10.90 | 3.33 | 10.61 | 0.05 - 16.14 |
| **Beta**<br>**(13 - 30 Hz)** | Smartphone | 5.22 | 2.34 | 4.94 | 0.88 - 11.15 |
|  | Passive Viewing | 6.07 | 2.30 | 5.99 | -0.21 - 11.00 |
| **Gamma**<br>**(30 - 45 Hz)** | Smartphone | 3.18 | 1.03 | 3.24 | 1.32 - 5.16 |
|  | Passive Viewing | 3.07 | 1.09 | 3.05 | 0.42 - 5.58 |

**Figure 4.**
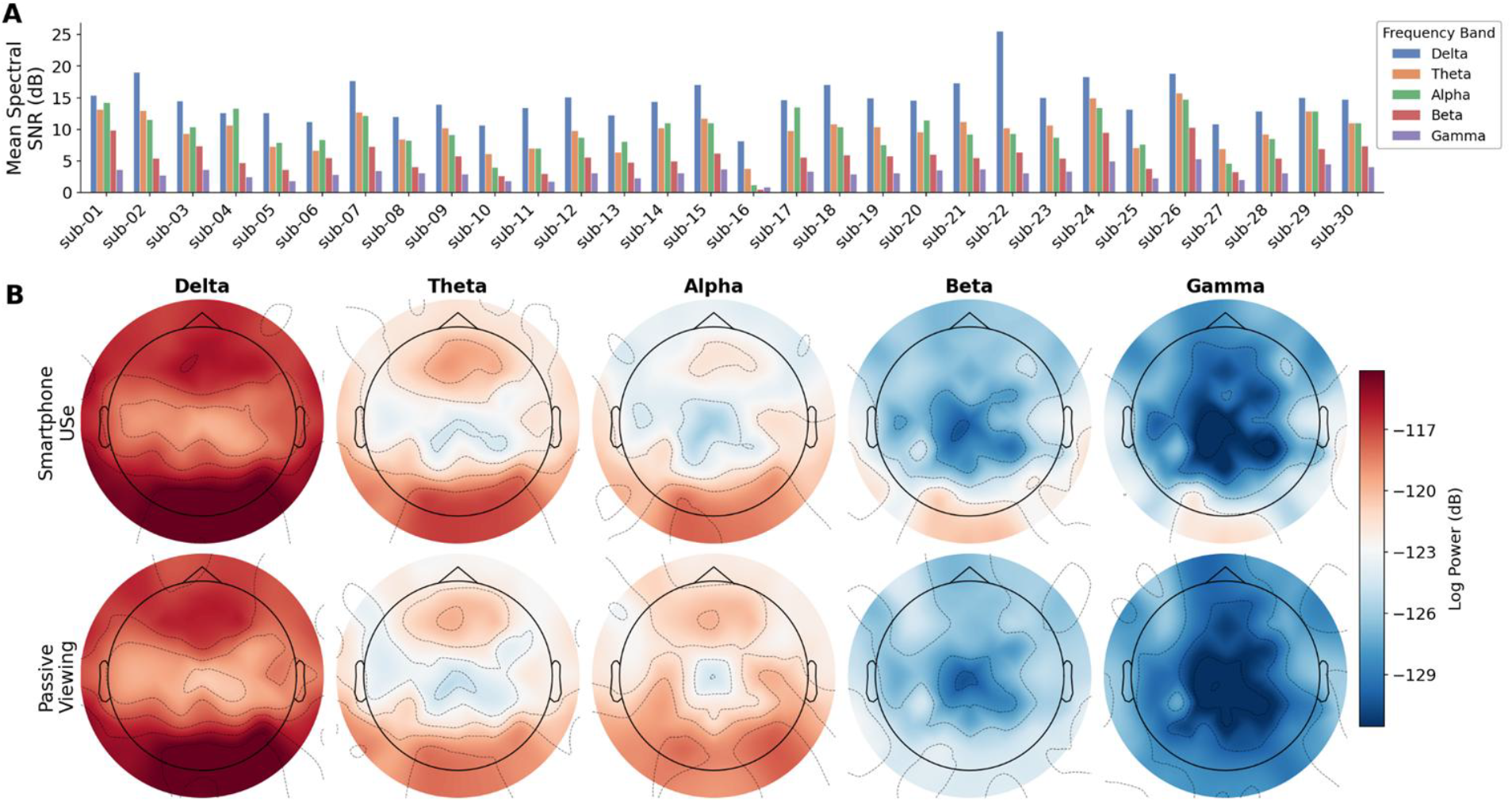
EEG data quality assessment using spectral signal-to-noise ratio and band-specific topographic power distributions. (A) Mean spectral signal-to-noise ratio (SNR) per subject for canonical EEG frequency bands (Delta, Theta, Alpha, Beta, and Gamma). SNR was estimated using Welch’s power spectral density and computed as the ratio of band-limited power to the high-frequency noise floor (45–95 Hz), expressed in decibels. (B) Grand-average scalp topographies illustrating the spatial distribution of log-transformed spectral power across conditions. Rows correspond to experimental conditions (Smartphone usage and Passive Viewing), while columns represent EEG frequency bands. Warmer colors indicate higher spectral power. The shared color scale allows direct comparison across bands and conditions. Topographies are computed using a 20% trimmed mean across participants to reduce the influence of extreme values.

Grand-average log-power topographic maps show physiologically plausible spatial distributions across all frequency bands in both conditions (Figure 4B). Delta power is strongest over frontal and vertex regions, theta showed a fronto-central distribution, and alpha demonstrated a clear posterior increase in the passive-viewing session compared to the smartphone usages session, consistent with known visual alpha effects. Beta and gamma bands show low, spatially diffuse power without focal temporal or frontal hotspots, suggesting minimal residual muscle artefact after preprocessing. Overall, the observed topographies are consistent with established EEG spatial patterns, supporting the technical validity of the dataset.

#### Eye-Tracking Data Quality and Oculomotor Features

Eye-tracking data quality is evaluated using a session-level processing pipeline that identifies experimental segments and computes behavioral metrics. Hardware TTL synchronization pulses (stream type = “sig”; direction = “in”; value = 1 (high)) are used to identify the onset and offset of each experimental session. Session boundaries are signaled by 5-pulse bursts at a nominal inter-pulse interval of 1 s (acceptance window: 0.95-1.05 s). Across all 30 participants, the extracted SM session durations spanned 609.85-609.91 s (mean = 609.90 s, SD = 0.01 s), demonstrating sub-second timing stability across the entire cohort. PV session durations ranged from 308.93 to 309.96 s (mean = 309.90 s, SD = 0.18 s).

Pupil diameter is filtered to the physiological range of 1.5-9.0 mm, retaining only samples with status = 0. All session means fell well within the accepted mesopic-photopic range (2.76-6.10 mm), confirming absence of systematic out-of-range contamination after filtering. Under typical physiological conditions, inter-ocular pupil diameter asymmetry (anisocoria)^43^ does not exceed 1 mm. Mean absolute bilateral difference is 0.19 mm in SM and 0.28 mm in PV, both well within this bound (right column; Figure 5). The maximum bilateral difference observed is 0.60 mm (sub-01, SM) and 0.78 mm (sub-02, PV), neither reaching the 1 mm clinical criterion. These values confirm acceptable tracker performance across both eyes for all participants.

**Figure 5:**
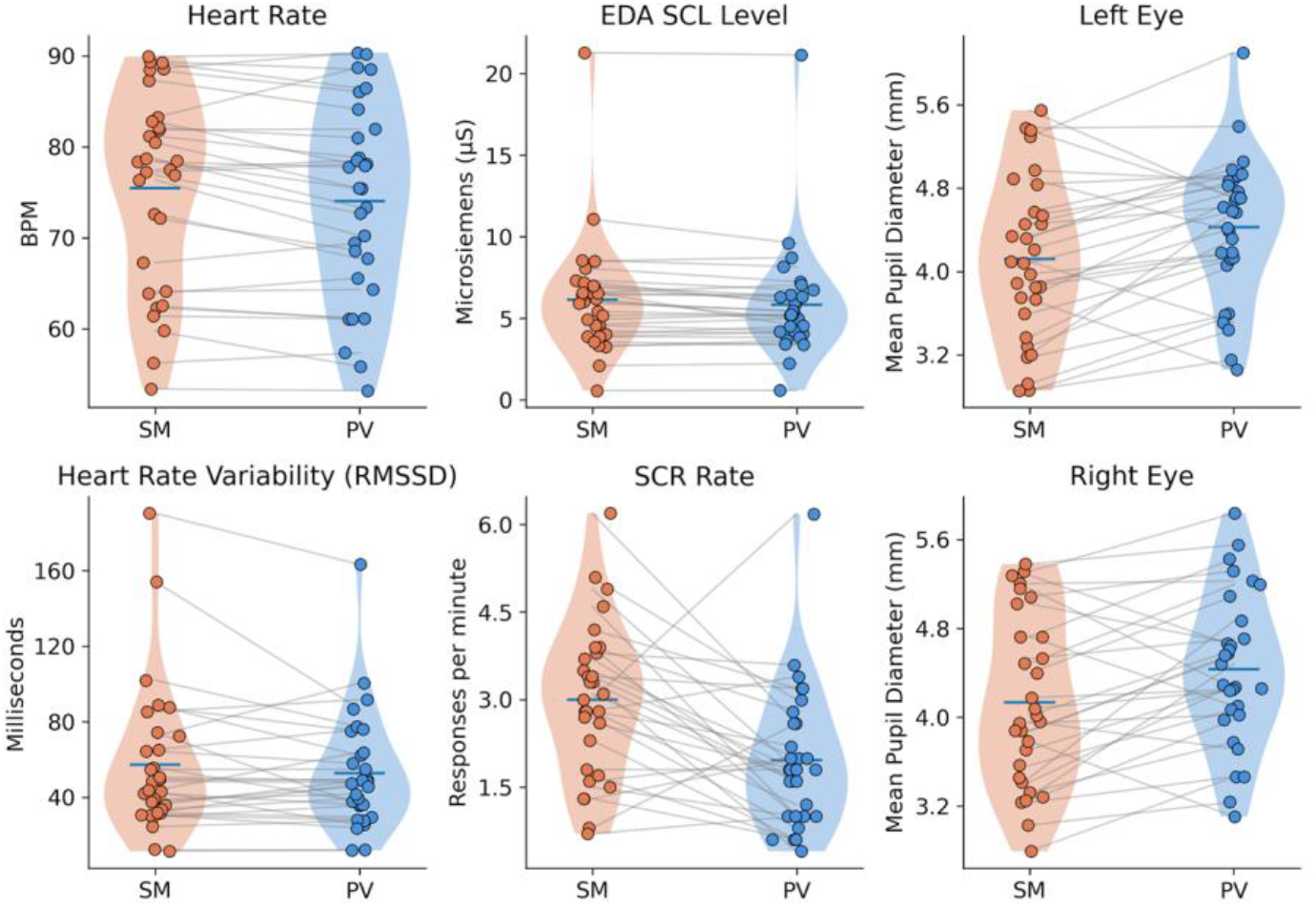
Subject-level physiological responses during Smartphone Use (SM) and Passive-Viewing (PV). Violin plots with overlaid individual participant data showing physiological metrics during the smartphone-use (SM) and passive-viewing (PV) conditions. Gray lines connect paired measurements from the same participant across conditions. Panels display heart rate (beats per minute), tonic electrodermal activity level (microsiemens), mean pupil diameter for the left and right eyes (mm), heart rate variability quantified by the root mean square of successive differences (RMSSD, milliseconds), and the rate of skin conductance responses (SCRs per minute). Violin distributions illustrate the variability across participants, with horizontal bars indicating the group mean for each condition.

Gaze coordinates are normalized to [0, 1] with the origin at the top-left corner of the display field. All gaze metrics are computed from status = 0 samples only. Grand-mean horizontal gaze is 0.528 (SD = 0.051) in SM and 0.493 (SD = 0.046) in PV, both consistent with central screen fixation (expected value ≈ 0.5). A large and condition-expected difference in mean vertical gaze is observed. SM sessions show a grand-mean vertical gaze of 0.570 (SD = 0.106), reflecting fixation directed at the lower-center screen region typical of smartphone use from a seated position, while PV session shows a substantially higher fixation position (mean vertical gaze = 0.315, SD = 0.096), corresponding to the upper portion of the screen. This downward-to-upward shift between conditions (Δ ≈ 0.25 normalized units) is physiologically and ecologically plausible and constitutes a positive validity indicator, as it reflects the distinct visual demands of sedentary computer use versus naturalistic video viewing posture^44^.

Head motion is captured by a triaxial accelerometer (stream type “ac”; units: m s^-2^) and triaxial gyroscope (stream type “gy”; units: ° s^-1^). Three-axis signals are reduced to Euclidean scalar magnitudes M = *√*(x^2^ + y^2^ + z^2^) using only status = 0 samples. During sedentary sitting, the accelerometer primarily captures gravitational acceleration (≈ 9.81 m s^-2^). Mean SM accelerometer magnitude across the cohort is 9.214 m s^-2^ (SD = 2.122), consistent with a combination of gravitational and minor postural-sway components. PV mean is 9.838 m s^-2^ (SD = 1.374), with virtually identical values across participants. Mean gyroscope magnitude is 1.631 ° s^-1^ (SD = 0.811) in SM and 1.382 ° s^-1^ (SD = 0.544) in PV. Contrary to the naïve expectation that freer body movement in PV would produce higher head rotation rates, 21 of 30 participants showed higher gyroscope magnitude during SM than PV. This pattern is ecologically plausible as during the SM condition, participants engaged in active smartphone tasks (typing, scrolling, reading) that involve frequent small head reorientations and postural adjustments, whereas the PV condition involved more passive, directed viewing with reduced head movement.

In addition to the quality metrics, the eye-tracking derivative dataset includes smartphone interaction annotations generated using Tobii Pro Lab. Interactions are manually annotated as *tap* (touchscreen tap), *swipe* (screen swipe gesture), and *drag* (touch-and-drag gesture), providing behavioral labels throughout the smartphone session with approximated temporal accuracy 0.5 sec. We note that the event annotations lack temporal precision, and are not suitable for event-related analyses. They may instead be used as a measure of interaction frequency over a longer timescale of several seconds To preserve participant privacy, application content and screen-level activity are not recorded; consequently, fine grain content-level annotations are not included in the dataset.

#### PPG and GSR Signal Validation

PPG and GSR signals are analyzed to evaluate the reliability and physiological plausibility of autonomic measurements during smartphone use and passive-viewing conditions. The PPG signal is band-pass filtered between 0.5-5 Hz to isolate the cardiac component while suppressing low-frequency drift and high-frequency noise. Systolic peaks are detected from the filtered waveform, and inter-beat intervals are computed to estimate heart rate and heart rate variability (HRV). Across the cohort (N = 30), all session means fall within the expected resting-to-light-activity range for ambulatory healthy adults. The cohort mean HR is 75.44 bpm (SD = 10.76) in SM and 74.02 bpm (SD = 10.79) in PV, a mean within-participant difference of 1.42 bpm (SD = 2.70).. HRV is quantified using the root mean square of successive differences (RMSSD), a time-domain metric reflecting parasympathetic modulation of cardiac activity, here derived from pulse-to-pulse intervals. Cohort RMSSD means (SM: 57.39 ms; PV: 52.78 ms) are consistent with published normative values for healthy adults during resting or passive conditions. The large within-cohort standard deviation (SM = 38.82; PV = 30.65) reflects normal physiological heterogeneity in vagal tone.

Electrodermal activity is decomposed into tonic (skin conductance level, SCL) and phasic (skin conductance response, SCR) components using the cvxEDA convex-optimization method^45^, after downsampling to 10 Hz. This model-based decomposition explicitly separates the slow tonic baseline from sparse phasic responses, which a single low-pass filter cannot achieve because the two components overlap in frequency. SCRs are detected on the phasic component using a minimum-amplitude criterion of 0.05 µS^46^, and SCR rate is expressed as qualifying responses per minute. Mean SCR rate is higher in SM (3.00 /min, SD = 1.31) than PV (1.96 /min, SD = 1.21), with 24 of 30 participants showing a lower rate in PV. Mean SCR amplitude is likewise greater in SM (0.34 µS, SD = 0.43) than PV (0.24 µS, SD = 0.30). Tonic EDA, computed as the mean of the recovered SCL component, is also higher in SM (6.13 µS, SD = 3.57) than PV (5.82 µS, SD = 3.44). SCL values across both sessions and all participants range from 0.568 to 21.283 µS, within the physiological range reported for healthy adult skin conductance. The convergence of three independent electrodermal measures, SCR rate, SCR amplitude, and tonic SC, on higher sympathetic activity during smartphone use indicates that the pipeline captures physiologically plausible, condition-sensitive autonomic modulation.

#### Smartphone Questionnaire Validation

The completeness and internal consistency of the questionnaire data are assessed following dataset curation. No missing questionnaire responses are identified for either the original 33-item SAS or the original 27-item MPPUS. The SAS demonstrates high internal consistency (Cronbach’s *α* = 0.928), whereas the MPPUS demonstrates acceptable internal consistency (Cronbach’s *α* = 0.741). The observed score ranges (SAS: 36-161; MPPUS: 71-201) fall within the theoretical ranges of the respective instruments (SAS: 33-198; MPPUS: 27-270), with no floor or ceiling effects observed. The mean ± standard deviation scores are 106.2 ± 26.8 for the SAS and 135.4 ± 34.2 for the MPPUS, indicating substantial inter-individual variability in smartphone use behaviours across participants.

#### Usage Notes

This dataset provides synchronized EEG, eye-tracking, PPG, and GSR recordings acquired during naturalistic smartphone use and a standardized passive-viewing condition, together with smartphone interaction annotations and validated behavioural questionnaire scores. All data are organized according to the Brain Imaging Data Structure (BIDS; version 1.11.1), valadated with BIDS validator (version 3.0.1) and can be read with standard BIDS-aware tooling, including MNE-Python and MNE-BIDS for the EEG recordings. Reusers should note that modalities are stored at their native sampling rates rather than resampled to a common grid: eye-tracking gaze, head-mounted IMU, and peripheral physiological channels are written as separate _physio files distinguished by the recording-<label> entity, each with its own JSON sidecar specifying sampling frequency, units, and column names. Channel names appear only in the “Columns” field of the sidecar; the TSV files themselves are headerless, per BIDS convention. Synchronization markers are provided in the accompanying _physioevents files, and cross-modal analyses should use these to align streams to a common time base before resampling.

To guide downstream reuse, we categorize supported analyses according to whether they primarily draw on within-subject temporal dynamics or across-subject feature variance, since these differ substantially in statistical power given the current cohort size. The dataset’s high-resolution, sub-second synchronized recordings and fine-grained interaction annotations (tap, swipe, drag, and hand use) provide many repeated observations per participant, giving within-subject analyses strong statistical power that is largely independent of sample size. Primary within-subject use cases include time-resolved cross-modal coupling – e.g., phase-amplitude or cross-correlation analyses testing whether central neural oscillations (frontal theta, motor beta, posterior alpha) and peripheral autonomic or oculomotor signals (GSR, HRV, pupil diameter) co-vary in ways consistent with theoretical constructs such as engagement, arousal, or attentional allocation; visuomotor and motor-cortical dynamics related to individual touch gestures, including hand-specific lateralization; and repeated-measures contrasts between naturalistic smartphone use and the passive viewing condition. We note that constructs such as “engagement” are inferred from patterns of physiological co-variation rather than measured against an independent ground truth, and results should be interpreted accordingly.

Primary use cases also include across-subject classification of broad activity categories. The expanded cohort includes an approximately balanced split between interactive gaming and passive short-form video consumption (n ≈ 15 each), sufficient to support supervised classification of app type from multimodal features using participant-level validation (e.g., leave-one-subject-out cross-validation, rather than random partitioning of time windows, which would conflate app-type discrimination with subject identity). Because app type reflects a self-selected, between-subjects grouping rather than a controlled manipulation, such classification robustly establishes *that* physiology discriminates activity type but cannot on its own disambiguate whether this reflects the differing cognitive and motor demands of the activities themselves, stable individual differences between people who prefer one activity over another, or both.

Analyses that depend on across-subject variance in high-dimensional feature spaces, without the benefit of repeated within-subject sampling, constitute exploratory reuse scenarios given the cohort size (N = 30). These include relating participant-level physiological and behavioral features to individual psychometric measures of problematic smartphone use (SAS, MPPUS), and more fine-grained content-level classification (e.g., distinguishing specific video genres or gameplay sub-states), which is additionally constrained by the absence of detailed content annotation beyond gesture type and hand use. Such analyses are well suited to hypothesis generation and methodological benchmarking but should not be interpreted as validated predictive or diagnostic models.

The naturalistic design introduces variability in application type, interaction behaviour and posture, device characteristics and viewing distance, and user settings; therefore, downstream analyses should account for these sources of heterogeneity when interpreting physiological or behavioural differences.

## Data availability

Dataset is available at OpenNeuro (*https://openneuro.org*) with accession number: ds007537 with doi:10.18112/openneuro.ds007537.v2.0.2.

## Code availability

Validation analysis was performed with Python and the corresponding code is available at https://github.com/csndl-iitd/ecf-natural-smartphone-neurophysiology

